# Identifying genes and pathways linking astrocyte regional specificity to Alzheimer’s disease susceptibility

**DOI:** 10.1101/2022.11.16.515390

**Authors:** Ran Zhang, Margarete Knudsen, Pedro Del Cioppo Vasques, Alicja Tadych, Patricia Rodriguez-Rodriguez, Paul Greengard, Jean-Pierre Roussarie, Ana Milosevic, Olga Troyanskaya

## Abstract

Astrocytes have been shown to play a central role in Alzheimer’s Disease (AD). However, the genes and biological pathways underlying disease manifestation are unknown, and it is unclear whether regional molecular differences among astrocytes contribute to regional specificity of disease. Here, we began to address these challenges with integrated experimental and computational approaches. We constructed a human astrocyte-specific functional gene network using Bayesian integration of a large compendium of human functional genomics data, as well as regional astrocyte gene expression profiles we generated in the mouse. This network identifies likely region-specific astrocyte pathways that operate in healthy brains. We leveraged our findings to compile genome-wide astrocyte-associated disease-gene predictions, employing a novel network-guided differential expression analysis (NetDIFF). We also used this data to predict a list of astrocyte-expressed genes mediating region-specific human disease, using a network-guided shortest path method (NetPATH). Both the network and our results are publicly available using an interactive web interface at http://astrocyte.princeton.edu. Our experimental and computational studies propose a strategy for disease gene and pathway prediction that may be applied to a host of human neurological disorders.

## Introduction

Alzheimer’s disease (AD), as with many neurological diseases and disorders, manifest in cell-type and brain-region-specific manner. During disease progression, AD is shown to present region-specific pathologies (1,2), with its first onset happening in selective neurons in the hippocampal region and has a clear region-specific progression. Besides, although AD has been characterized by impaired neurons, the role of glial cells has been increasingly recognized (3–6). Understanding the cell-type and region-specific regulation of AD pathology is important for discovery of new AD risk genes and development of targeted treatments on malfunctions in selected cell types/regions.

Astrocytes, the most abundant glial subtype, have a major role in supporting healthy neuronal function. That includes synaptic, trophic, structural, and metabolic support, removal of oxidants and maintenance of adequate extracellular ion concentrations, water exchange, and the regulation of blood-brain barrier permeability and function (7,8). Astrocytes are a morphologically and molecularly heterogeneous cell type in respect to their spatial distribution, as well as developmental and postnatal period (6,8–11).

Astrocytes have been implicated in many psychiatric and neurodegenerative diseases, including AD. However, the molecular machinery mediating astrocytes’ connection to AD is not fully understood, and it is unclear if astrocytes contribute to the increased vulnerability presented by certain brain regions in AD. While Genome-wide association studies (GWASes) are successful at identifying disease genes, they unfortunately do not provide information regarding the cell type mediating the effect of a disease gene, or the tissue/region in which these effects will be strongest. Besides, the majority of the genes captured in neurological disease GWAS are not astrocyte-specific (6,12,13), implying that astrocytes might not harbor the main genetic cause, yet they might be important disease modulators, and potentially important targets for therapeutic interventions.

To understand astrocytes’ role in AD pathology, we must be able to systematically investigate astrocytes’ region-specific functions and map the molecular machinery that is dysregulated within astrocytes in specific brain regions. The challenge here is to extract biological meaningful information from the marginally different astrocytes profiles across regions, to discover complex molecular connections between these regional astrocyte regulations and AD, and to transfer the high-confidence experimental knowledge that are usually derived in mouse to human pathology. Currently, studies on regional specificity of astrocytes rely on the comparison of astrocyte transcriptomes for the identification of individual differentially expressed genes between regions (14–17). In the context of AD, most recent studies are focused on describing the gene expression state of astrocytes in disease context (18–21) and identified genes and pathways that are perturbed in AD, they do not provide regional specificity and fall short of mapping the complex interactions between region-specific gene expression, pathways, and human AD pathology. Additionally, the role astrocytes play in disease may not be revealed from differential expression studies alone, as cellular functions may be disrupted through network effects that are not evident from transcriptional analysis.

Here we propose a systems-level integrative framework to investigate the role of astrocytes in AD with a goal of dissecting region-specific molecular machinery in astrocytes that contributes to AD pathology. To do that, we first integrate 276 brain -omics datasets in an astrocyte specific context by using high resolution *ex vivo* regional astrocyte data from mice to build a human astrocyte gene-gene functional interaction network. Integrating this human astrocyte network with AD quantitative genetics data, we then develop several novel methods to identify hippocampus-specific AD signatures in astrocytes and discover mediator genes and pathways through which hippocampal astrocytes contribute to AD pathology. We provide an interactive web interface to the human astrocyte network and the resulting findings at http://astrocyte.princeton.edu.

## Results

### Modeling functional relationships of genes in human astrocytes with an integrative network approach

To characterize region-specific astrocytes, we first experimentally generated astrocyte gene expression profiles in five major brain regions in mouse: prefrontal cortex (P), hippocampus (H), amygdala (A), caudate nucleus (C) and ventral striatum (V) (Figure 1A, Methods). For this purpose we utilized TRAP-seq (22), a method that enables detailed profiling of the actively translated rather than just transcribed mRNA, thus better capturing the functional state of the cell, enabling us to generate *ex vivo* region-specific astrocyte signatures.

**Figure 1:**
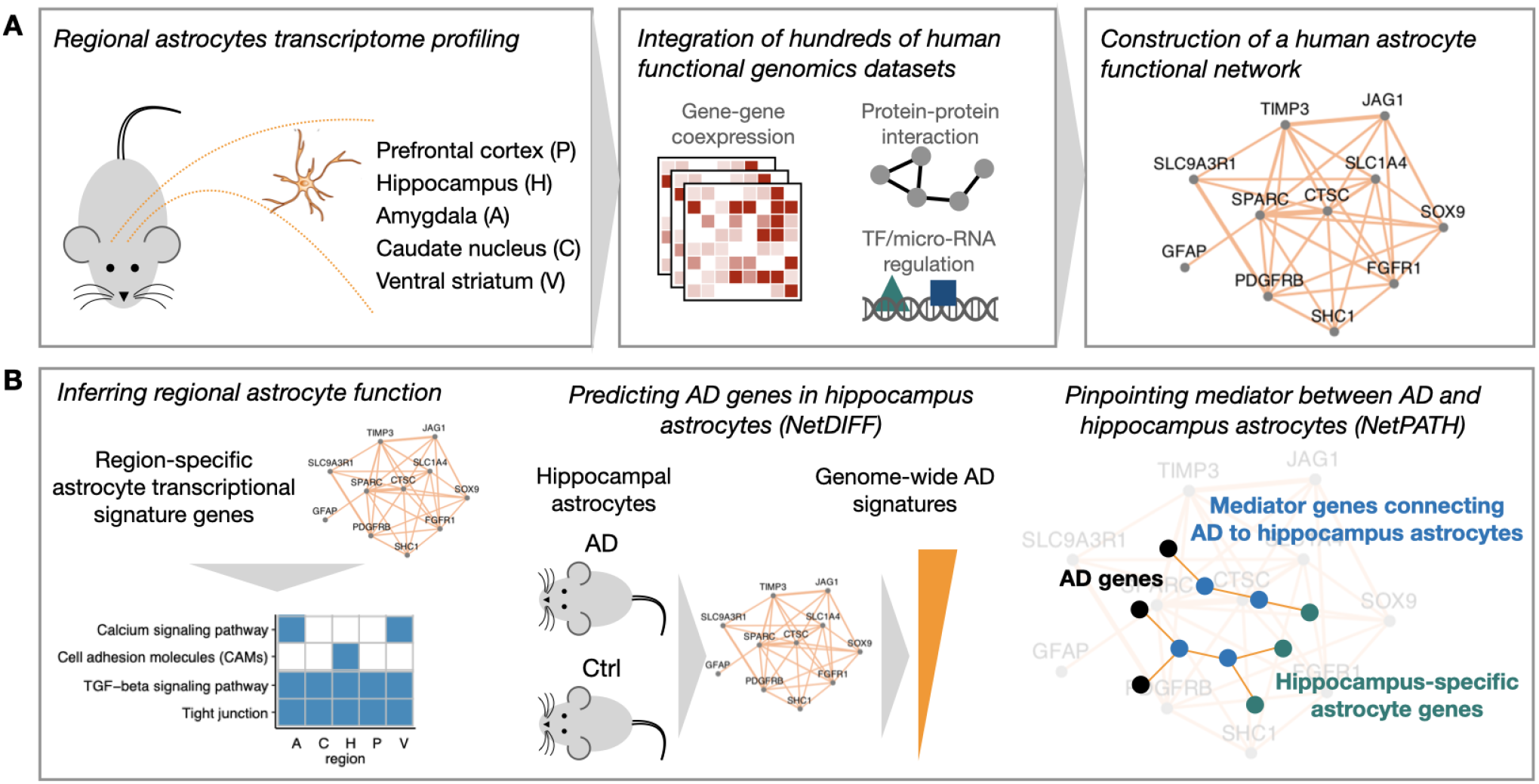
Overview of the study - construction and applications of astrocyte-specific networks. **A:** astrocyte-specific functional network integration. Left: astrocyte signatures were extracted from the regional astrocyte gene expression profiles; Middle: the signatures were then used to construct a human astrocyte specific functional network through Bayesian integration of human brain functional genomic datasets. Right: An illustrative example of a small part of the resulting network, each node is a gene and edge weight is the likelihood of a pair of genes to participate in the same pathway/biological processes in astrocytes. **B:** downstream applications. The astrocyte network guides prediction of region-specific pathways in astrocytes (left), helps pinpoint regional disease-relevant genes from noisy disease models (middle), and facilitates reprioritization of novel astrocyte disease-associated genes (right).

In order to identify the region-specific astrocyte functions and association with AD pathology in human, we leverage these high-resolution mouse signatures and human brain expression data to extract meaningful regional and disease signals from subtle differences across regional astrocytes. We first constructed a general human astrocyte functional gene interaction network to systematically capture the overall gene relationships in human astrocytes (Figure 1A). The network is built through regularized Bayesian integration that leverages the high confidence astrocytes signatures generated in mouse experiment and large scale human functional genomics datasets in the brain, to capture functional gene relationships in human astrocytes (Figure 1A, Methods) (23). Intuitively, the mouse data provides region-specific signals that are combined with experimentally-validated Gene Ontology biological pathways in a probabilistic framework that extracts astrocyte-relevant signals from each human brain dataset and generates a human astrocyte network. In this network, each node is a gene and the edge weight between two nodes represents the strength of the functional relationship between that gene pair. Then, we use this robust network representation of astrocytes as a foundation for downstream studies, proposing several algorithms to dissect the roles of regional astrocytes in AD pathology (Figure 1B).

### Region-specific astrocyte gene signatures and functions

To identify functional heterogeneity of astrocytes based on their anatomical location, we compared astrocyte profiles obtained from the five above-mentioned brain regions. A multidimensional scaling (MDS) plot (Figure 2A) shows that astrocytes TRAP-seq profiles cluster by region, indicating the existence of regional heterogeneity in astrocyte populations. One exception was that the astrocytes from ventral striatum and caudate nucleus clustered together, suggesting that in the striatum subregional differences in gene expression are not enough to separate these samples.

**Figure 2:**
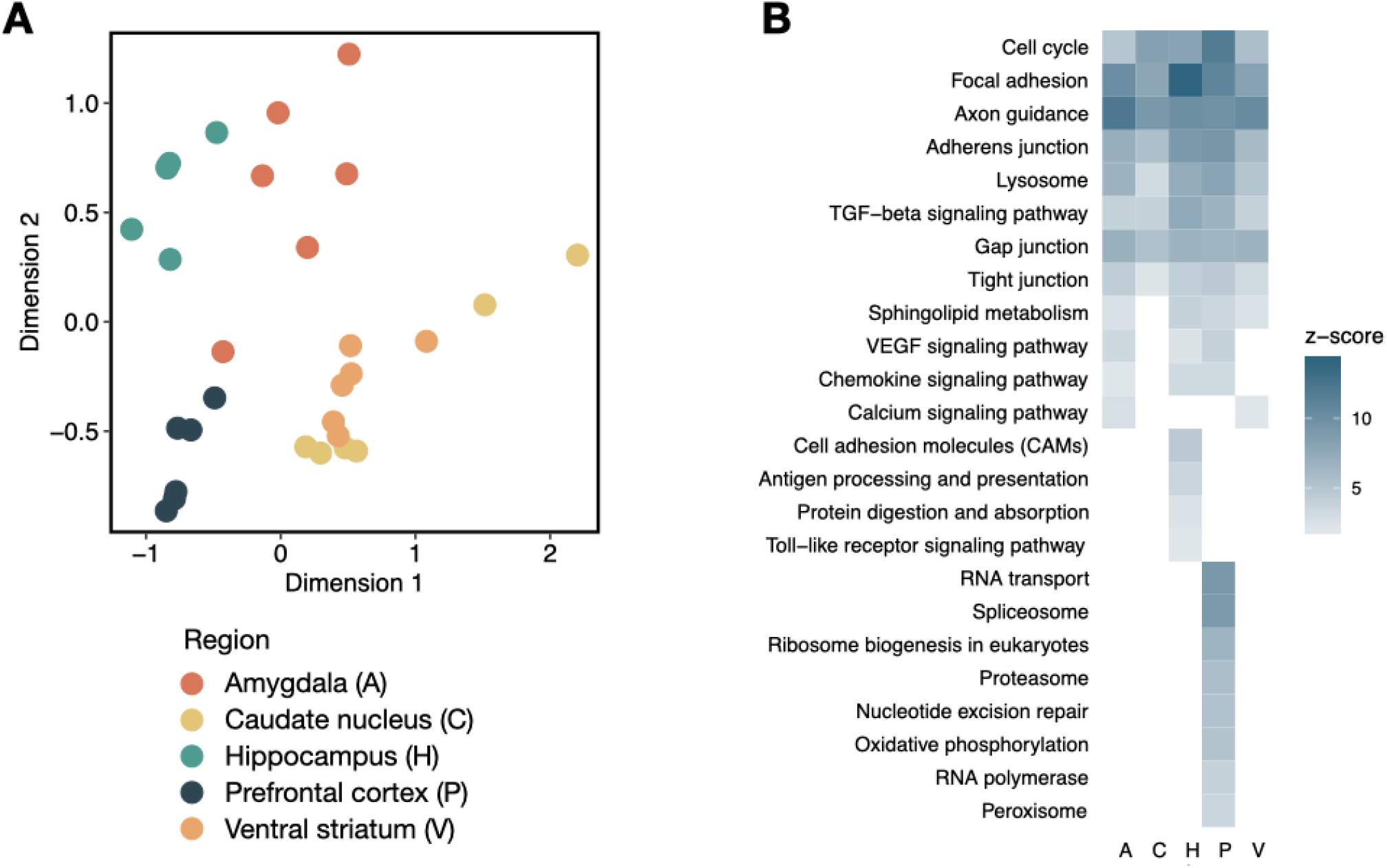
Region-specific astrocytes transcriptomic profiles and pathway enrichment. (A) Multidimensional scaling plot of Trap-seq profiles of individual regional astrocyte samples suggests that astrocyte transcriptomic profiles cluster by region. Each sample is represented as a dot and colored by region. (B) Heatmap of KEGG pathways involved in region-specific astrocyte regulation. KEGG pathways are identified through rank-based enrichment tests on region-specific astrocyte functional gene signatures and colored by enrichment z-score (pathways with FDR≤0.05 are colored in blue). Only selected pathways are plotted for visual clarity.

We then defined region-specific astrocyte transcriptional signatures to include all genes that are significantly higher expressed in one region compared to other regions (FDR<=0.01, CPM>=10, Methods). These signatures represent the most characteristic transcriptional molecular features of astrocytes within each brain region, and thus were used to identify region-specific biological pathways.

However, since the signatures were generated based on differential gene expression, they are likely to represent only part of the functional differences between astrocytes from different regions. For instance, genes coding for proteins that are differentially phosphorylated, or physically interacting with proteins coded by region-specific gene signatures, will not be captured by these signatures. To address this challenge and focus on region-specific astrocyte signals relevant to the human brain, we derive region-specific astrocyte functional signatures by using the identifying genes that are functionally connected to the transcriptional signature genes in the human astrocyte functional network.

We then identified the most salient processes for astrocytes in different brain areas by rank-based KEGG pathway enrichment analysis of region-specific astrocyte functional signature genes (Figure 2B). Our analysis indicates that while some biological pathways are shared across regions, some represent region-specific functions (Figure 2B). In general, we observe more region-specific pathways in prefrontal cortex and hippocampus regions, suggesting that astrocytes in these regions might have been adapted to have a more specific role. For example, cell adhesion molecules are enriched in the hippocampus, which may point to a greater need for cell-cell communication and organization in the hippocampus because of the large number of complex circuitry connections within the structure and with other brain regions. This also suggests hippocampal astrocytes might have particularly specific high levels of plasticity and cell adhesion underlying both development and maintenance of synaptic function in the hippocampus (8,24). As another example, peroxisome, RNA and proteasome related pathways are predominantly enriched in the prefrontal cortex. Prefrontal cortex is involved in cognition, decision making, and reward regulation (25–28) potentially suggesting the need for faster, more efficient metabolic pathways and clearance of the free radicals and metabolic byproducts. On the other hand, there are a number of pathways and genes that appear in every region assayed, alluding that they may have a more expansive, broader role in the brain, such as regulation of actin cytoskeleton and lysosomes. Interestingly, a large number of these pathways are essential during development, such as axonal guidance; while other processes, such as tight junction, adherens junction, gap junctions, and focal adhesion, are critical for communication both within astrocytes and between astrocytes and other cells.

### A novel framework predicts AD signatures in region-specific astrocytes

AD is a neurodegenerative disease with a clear region-specific manifestation. Pathological lesions form in the hippocampus, where astrocytes are thought to play a role (29), at earlier stages of AD than the other regions assayed here. With the finding of regional heterogeneity in astrocytes, we further hypothesized that hippocampus-specific features of astrocytes may contribute to this region-specific AD pathogenesis.

To understand regional astrocyte’s role in hippocampus AD pathogenesis, we profiled hippocampal astrocytes in healthy mice and the 5XFAD AD mouse model at 14 weeks of age (Figure 3A, Methods). Differential expression analysis between AD and control transcriptional profiles were only able to identify 33 genes significantly up- or down-regulated in AD (FDR<=0.05, Table S4). To focus on human-relevant signals and mitigate the limitations in differential expression to identify disease-associated genes, we developed a novel NetDIFF method. In NetDIFF, support vector machine classifiers were trained with the astrocyte network as the input, and top differentially expressed genes in the mouse study serving as positives and non-differentially expressed genes as negatives. Intuitively, this identifies astrocyte network patterns preferentially associated with differentially-expressed genes and then identifying other disease-associated gene candidates based on similar network connectivity. NetDIFF therefore reprioritized candidate disease genes based on integrating differential gene expression patterns and functional gene interaction patterns in the human astrocyte network (reprioritized genes in table S4).

**Figure 3:**
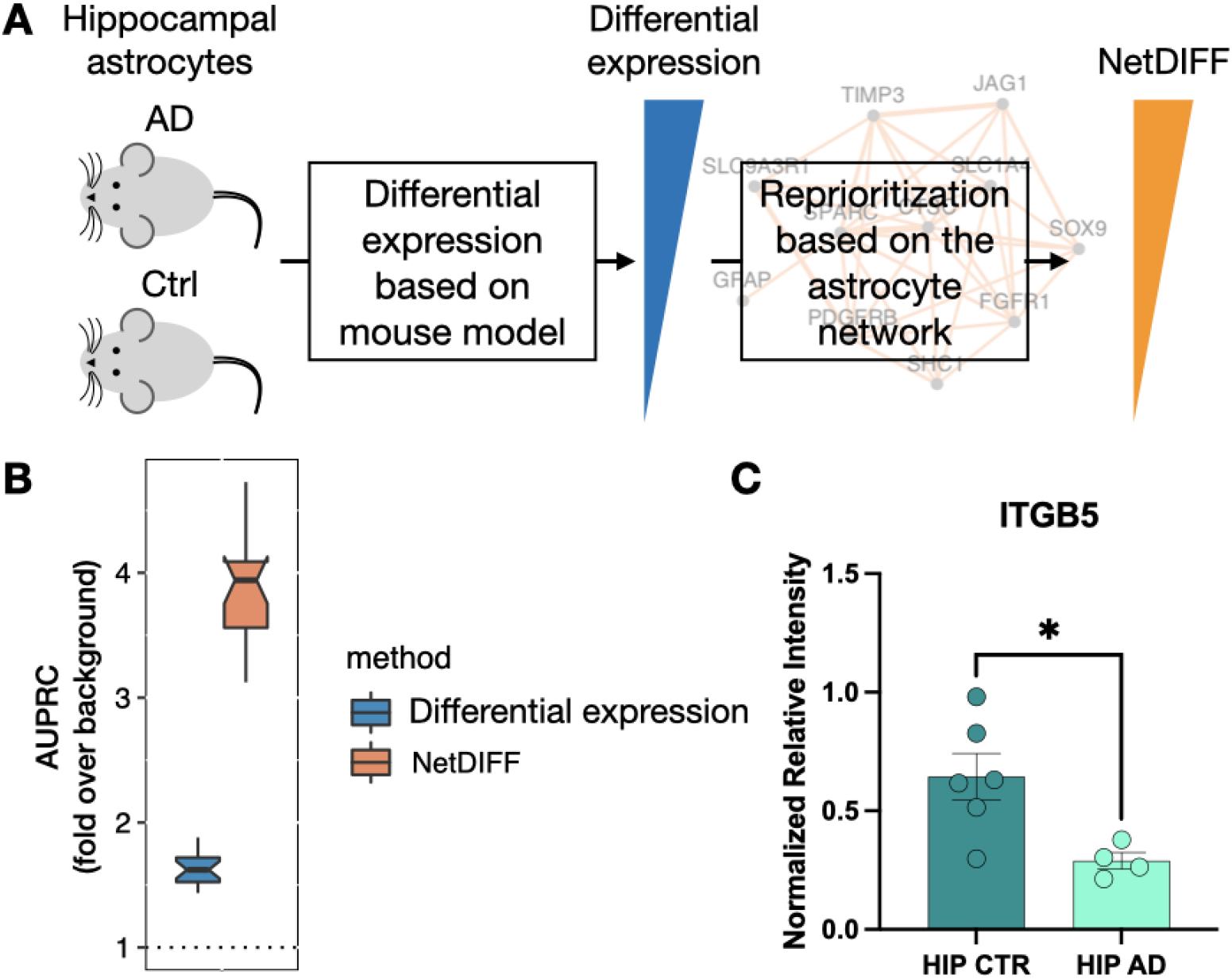
Astrocyte disease signatures prediction with NetDIFF. **(A)** Illustration of NetDIFF algorithm. We first calculated a differential gene expression pattern based on AD v.s. Control mice. The NetDIFF algorithm then reprioritized differentially expressed genes based on the functional gene interaction pattern in the network to generate NetDIFF predictions. **(B)** NetDIFF predictions significantly outperformed original differential expression analysis in astrocyte AD gene prediction. The predictions were evaluated on independently expert-curated astrocyte AD genes. Background performance was generated by training on permuted gold-standards (Methods). **(C)** Western blot quantification of ITGB5 protein expression in AD vs. control human brain slices. The top prediction, ITGB5, is significantly upregulated at protein level in human AD vs. control hippocampus on western blot.

To test if NetDIFF better captures AD genes than differential expression analysis alone, we systematically evaluated the predictions on an independent expert-curated astrocyte AD gene set. This set included genes with demonstrated functional association with AD pathology (e.g. amyloid-β and memory) in previous studies (see curation criteria in the methods). NetDIFF is significantly better at identifying known curated AD genes than differential expression alone (Figure 3B), indicating that NetDIFF could facilitate disease gene discovery with improved accuracy compared to differential expression analysis.

The top NetDIFF predicted gene, integrin subunit beta 5 (ITGB5), assembles with ITGV integrin subunits into ITGB5/ITGV heterodimers on the cell membrane, where it is involved in cell-surface mediated signaling (30,31). Reduced protein levels of ITGB5/ITGV have been found in the plasma of patients with cognitive impairment or dementia (32). In our study, ITGB5 gene is not significantly differentially expressed between healthy and AD mice in the hippocampal astrocytes, but based on the combination of its differential expression and network connectivity, NetDIFF reprioritized it as the top AD signature gene in hippocampal astrocytes. We thus hypothesized that ITGB5 may be differentially expressed at protein level in the hippocampus of human AD patients vs controls. We experimentally tested this hypothesis by performing western blot using human hippocampal samples. We also assessed the regional specificity of this interaction by performing the same experiment in the frontal cortex samples. Indeed, we found that the ITGB5 protein level is significantly reduced (p-value=0.021) in the hippocampi of AD patients compared to control (Fig. 3C). This demonstrates that NetDIFF can identify novel candidate disease genes in human.

### Predicting region-specific mediators of AD manifestation

We then aimed to pinpoint how the genetic causes of AD contribute to region-specific astrocyte dysfunction by identifying genes mediating the connection between region-specific astrocyte function and AD pathogenesis. To address this challenge, we developed NetPATH, an approach for identifying genes that act as mediators that connect known AD genes to region-specific astrocyte transcriptional signature genes. Intuitively, NetPATH uses the human astrocyte functional network to identify genes that are most critical to the shortest paths connecting known genetic causes of AD with astrocyte transcriptional signature genes from the hippocampus (Methods).

We applied the NetPATH method to investigate the connection of astrocytes in the hippocampus to genetic causes of AD, and identified 51 genes that are specifically connecting AD genetic risk genes hippocampus-specific astrocyte transcriptional signature genes, while not meditating other regional astrocyte’s connection to AD (see Methods). These genes are enriched in biological processes related to glial function and AD (Figure 4B).

**Figure 4:**
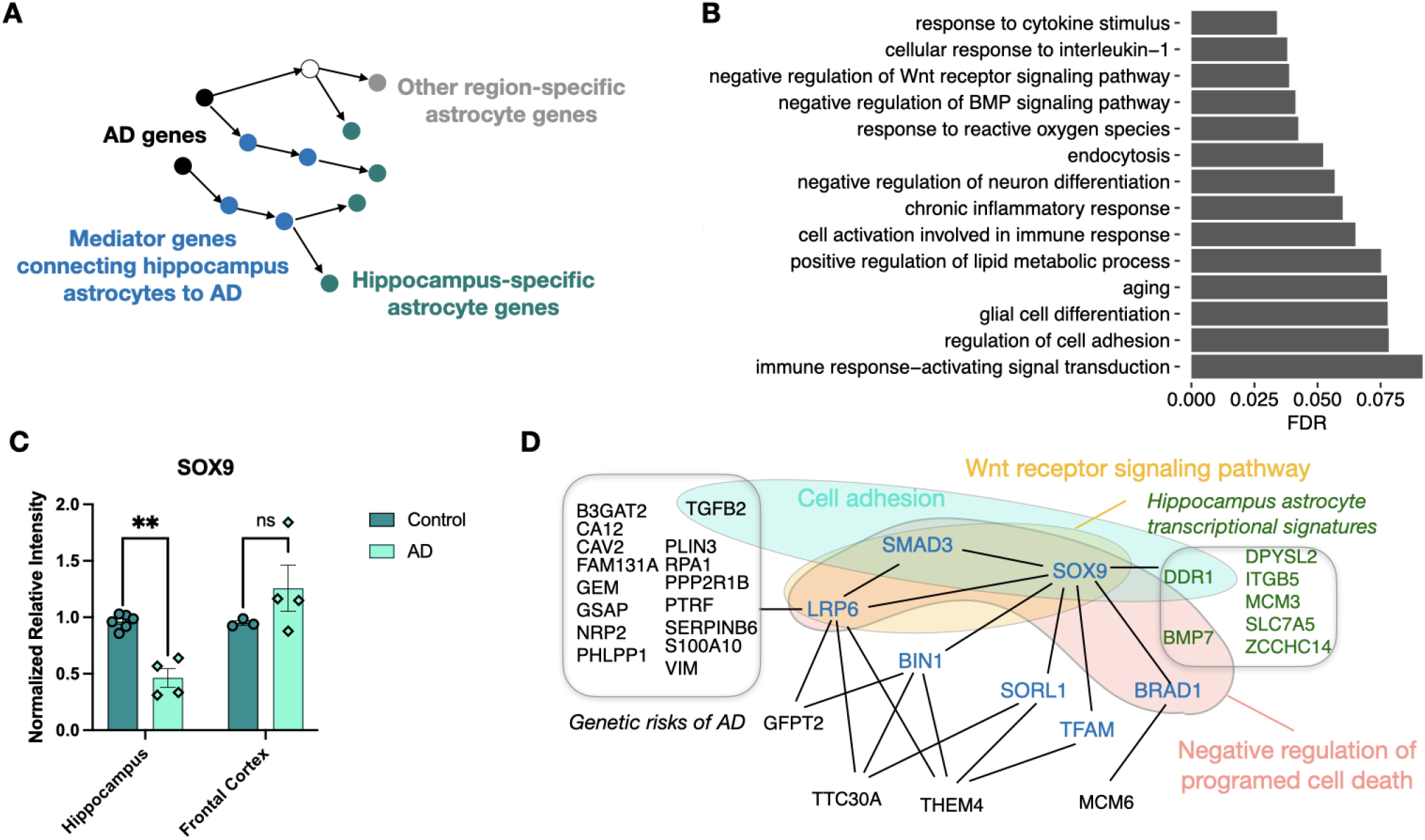
Identifying region-specific disease mediators using NetPATH. **(A)** Illustration of NetPATH algorithm. In the astrocyte network, we first identified shortest paths connecting AD genes (black) and hippocampus-specific astrocyte transcriptional signature genes (green). Genes enriched in these shortest paths compared to random permutations were retained as mediator genes (blue). **(B)** Top biological processes enriched in the hippocampus-specific AD mediators. **(C)** SOX9 is differentially expressed between AD and Control samples in the human hippocampus but not in the frontal cortex based on Western blot. **(D)** NetPATH-identified SOX9 associated network mediators of hippocampal astrocyte connection to AD genetics. NetPATH identified mediators (blue) and biological processes (colored shapes) mediating the connection between AD genes (black) and hippocampus astrocyte transcriptional signature genes (green) (only SOX9 mediation relevant AD genes and signature genes are shown).

One of the NetPATH-identified genes is SOX9, which is also a top-ranked NetDIFF prediction, suggesting it may be an important mediator of AD in human astrocytes. It has been shown that amyloid-β regulates expression of SOX9 specifically in astrocytes *in vitro (33)*. This suggests that SOX9 protein expression may be perturbed in human AD vs. healthy control astrocytes specifically in the hippocampus region. We found that SOX9 protein level was indeed significantly reduced in the hippocampi of the AD patients (p-value=0.007), while no significant change was detected in the frontal cortex between AD and healthy controls (p-value=0.164, Fig. 4C). SOX9-associated mediators between genetic AD risk factors and AD effects in hippocampal astrocytes appear to contain multiple members of the Wnt/beta-catenin signaling pathway (LRP6, SMAD3, SOX9) (Figure 4D). It has been shown that SOX9 regulates LRP6 and Wnt in other tissues (34,35) and neuron-specific lack of LRP6 leads to age-dependent loss of synaptic function and decline in cognition (36). Wnt signaling is dampened in the aging brain, leading to reduced synaptic plasticity and synaptic function (37), possibly by regulating microglia proliferation and survival (38). Here, our computational analysis identified astrocyte- and hippocampus-specific protein expression change in SOX9 (which we experimentally verified) and a major link connecting this change with the LRP6-Wnt signaling cascade in AD hippocampal pathology.

## Discussion

Traditional studies on regional astrocytes are mostly focused on region-specific transcriptomic profiles, and it is unclear whether the diversity of astrocytes across different brain regions corresponds to region-specific pathology of AD. To understand regional astrocyte regulation and association with AD pathology, here we developed a framework to extract the molecular dysfunctions and interactions of AD in region-specific astrocytes, by integrating our high-quality, targeted TRAP-seq experiments from mouse brain regions with large human brain data compendia from public databases to construct a human astrocyte functional network and identify the region-specific functions of astrocytes. We then generated an AD-specific TRAP-seq dataset from mouse hippocampus. We developed a machine learning based method (NetDIFF) to use these data and the human astrocyte network to prioritize genome-wide AD signatures in hippocampal astrocytes and a graph-based approach (NetPATH) to pinpoint hippocampal astrocyte gene mediators of AD region-specific pathology.

The regional diversity in astrocyte transcriptome and function raises a possibility that mutations or pathways dysregulation in astrocytes could have region-specific effects, which through cell-cell communication could contribute to selective vulnerability of neurons. In this study, we found region-specific astrocyte transcriptional signature genes shared cohesive roles in relevant pathways. For example, we found cell adhesion pathway genes to be enriched in the hippocampal astrocytes.

Hippocampus, with its complex circuitry (39,40) and high level of plasticity (41), may require tightly controlled synaptogenesis and synaptic activity. The astrocyte-specific cell adhesion molecules, with role in synaptogenesis and synaptic maintenance (8,24), provide this region-specific need for tighter control of homeostatic synaptic function.

We discovered ITGB5 as the top ranking hippocampal AD-associated gene in astrocytes and confirmed its depleted protein levels in the hippocampus of the AD patients. The only previously published data linking ITGB5 to AD was a biomarker study where increased protein levels in the plasma have been associated with a lower amyloid burden, and decreased odds of dementia and cognitive decline (32).

The cell type responsible and the causal relationship between ITGB5 and lower cognitive decline were not known. Here we found that ITGB5’s dysregulation in hippocampal astrocytes may play an important role in AD pathology, potentially mediated through SOX9 and Wnt signaling - a hypothesis that warrants further investigation. Specifically, lower expression of SOX9 in AD may disrupt the ITGB5 and LRP6 expression, leading to detrimental changes in cell adhesion, Wnt signaling, and cell death. It is also worth noting that reactive astrocytes do not appear to express SOX9, at least not consistently. Upregulation of (6), or downregulation (41), or no change (42,43) of Sox9 have been reported, depending on the way that reactive phenotype was induced, while ITGB5 is decreased in reactive astrocytosis induced by TNFa (42). One possible explanation for this is that the transition of astrocytes from homeostatic to reactive state is not synchronized between different subregions of the human hippocampus (11), and the expression level is dependent on the area used for the analysis.

Our study suggests that astrocyte’s region-specificity has an impact on AD neuropathology, and suggests several important questions for further investigation. First, it is still unclear whether astrocytes are responding to changes in their environment in AD pathology (such as amyloid-β, tau, TGFB and hallmarks of advanced AD), or are they drivers of such pathological changes through regulation of cell differentiation and synaptogenesis. In addition, the timeline of astrocyte-induced pathological changes during AD onset remains obscure. Furthermore, since astrocytes act in a supporting role, their interactions with other cell types, such as neurons and microglia, are critical for our understanding of regional AD pathology, which would be interesting to investigate in future.

In summary, we presented novel computational algorithms to integrate region-specific and AD mouse models, human functional genomics data from the public database, as well as the genetic risks of AD, to pinpoint the region-specific regulation of astrocytes and its hippocampus-specific role in human AD onset. We identified SOX9 and ITGB5 as genes that might contribute and mediate the hippocampus-specific role of astrocytes in AD, and validated their protein level expression change in human AD samples. Furthermore, our systematic approach provided several lists of gene and interaction candidates for further investigation. We also built an interactive web interface at http://astrocyte.princeton.edu for biologists to explore the results and generate their own hypotheses.

## Acknowledgements

The authors would like to thank Dr. Shai Shaham for thoughtful comments on the manuscript. We would also like to thank Drs. Nathaniel Heintz and Joseph D. Dougherty for the Aldh1l1-TRAP mouse line.

## Methods

### Mouse models

#### Regional astrocytes from wildtype mice

Mouse transgenic line Adlh1l1-TRAP was used for this project. Twelve to fourteen-week-old healthy male and female mice were lightly anesthetized with CO_2_ and the brains were quickly removed and dissected. Prefrontal cortex, caudate nucleus, ventral striatum, hippocampus and amygdala were dissected. For each brain region, three technical replicates were processed to gather the astrocyte-specific mRNA for sequencing. Technical replicates contained two pooled biological replicates for prefrontal cortex, caudate nucleus, and hippocampus, and six biological replicate samples for ventral striatum and amygdala.

#### Alzheimer’s disease model

For Alzheimer’s disease mouse model, we utilized two mouse lines, Aldh1l1-TRAP and 5xFAD (B6.Cg-Tg(APPSwFlLon,PSEN1*M146L*L286V)6799Vas/Mmjax; The Jackson Laboratory, strain # 008730). We used the 5xFAD Alzheimer’s disease model because it provided fast and prominent gliosis, enabling isolation of ample mRNA from TRAP mice at an earlier time. Control (Aldh1l1-TRAP only) and 5xFAD (Aldh1l1-TRAP x 5xFAD) mice were euthanized at 14 weeks of age and RNA was isolated from the whole hippocampus.

All mice were kept in the Rockefeller University animal facility with access to food and water *ad libitum*, in tempered animal rooms with a 12 hour light/dark cycle. All experimental procedures were approved by the Rockefeller University IACUC.

### Human tissue samples

The complete list of human tissue samples used for this study are in Table 1. Human fresh frozen tissue samples were obtained in collaboration with the Karolinska Institutet (Stockholm, Sweden). These included hippocampal and frontal cortex tissue of Alzheimer patients and age matched controls. All experiments involving the human brain tissue had appropriate IRB protocols approved before experiments commenced. All tissue samples were processed in accordance with the ethical and regulatory rules guiding the use of human samples for research.

**Table 1:** human tissue samples.

|  | sample ID | sex | age | post mortem/hrs | ApoE genotype | frontal cortex | hippocampus |
| --- | --- | --- | --- | --- | --- | --- | --- |
| CONTROL | 151 | N/A | 82 | 9 | E3/E3 | yes | yes |
|  | 172 | F | 80 | 7 | E3/E3 | yes | yes |
|  | 34 | F | 83 | 9 | E3/E3 | yes | yes |
|  | 82 | N/A | 69 | 22 | E3/E3 |  | yes |
|  | 85 | N/A | 71 | 12 | E3/E3 |  | yes |
|  | 75 | N/A | 56 | 19 | E3/E3 |  | yes |
| ALZHEIMER | 143 | N/A | 76 | 4 | E3/E4 | yes | yes |
|  | 86 | N/A | 82 | 24 | E3/E3 |  | yes |
|  | 173 | N/A | 98 | 31 | E3/E3 |  | yes |
|  | 126 | N/A | 75 | 24 | E3/E3 |  | yes |
|  | 96 | F | 89 | 34 | E4/E4 | yes |  |
|  | 182 | M | 74 | 12 | E4/E4 | yes |  |
|  | 192 | F | 83 | 8 | E4/E4 | yes |  |

### Immunoblot Analysis

Human hippocampal and frontal cortex extracts from Alzheimer’s disease patients and age-matched controls (list is shown in Table 1) were weighed and mechanically homogenized (20% weight/volume) in lysis buffer [1.5% SDS (w/v) in PBS with added EDTA-free protease inhibitor (Roche #11836153001) and PhosSTOP phosphatase inhibitor (Roche #4906845001)]. A bicinchoninic acid assay was used to determine the total protein concentration of samples, which were then set to 2 μg/mL per sample by adding more lysis buffer and equal amounts of 4x LDS sample buffer (Invitrogen #NP0007) with added 5% (v/v) ß-mercaptoethanol. 20 μg of protein from each sample were loaded in separate wells in a 4-12% Bis-Tris gel (Invitrogen #WG1403A) and ran at 90V using MOPS SDS running buffer (Invitrogen #NP0001), and later transferred to a PVDF membrane. Blot was then incubated in primary antibodies overnight at 4°C: anti-SOX9 (1:100, Santa Cruz # sc-166505), anti-ITGB5 (1:500, Santa Cruz # sc-398214), and anti-ß-Actin (1:10,000, Cell Signaling # 8H10D10). After incubation, blot was washed with TBST (0.2% Tween-20) and incubated with dye-conjugated secondary antibodies for 45 minutes at room temperature. Blot signals were recorded by Odyssey infrared imaging systems (LiCOR Biosciences), and analyzed using ImageJ. Protein levels were normalized to actin levels, and analysis was done using a two-way t-test in Prism.

### BacTRAP immunoprecipitation and mRNA isolation

We used the TRAP mouse line with the astrocyte-specific promoter for the gene for aldehyde dehydrogenase 1 like 1 (Aldh1l1) regulating the expression of L10a-GFP. Polysome immunoprecipitations were performed as described previously (22). Immunoprecipitation was performed overnight with the mixture of two GFP monoclonal antibodies. mRNAs were purified using Qiagen Rneasy Micro Plus kit, following the manufacturer’s protocol, and quantity and quality were determined with a Nanodrop Spectrophotometer and Agilent Bioanalyser. For each sample, mRNAs were amplified using the Ovation RNA-seq System V2 (NuGEN). Library preparation and amplification were done by the Rockefeller University Genomic Facility, with Illumina HiSeq platform used for sequencing.

### RNA-seq data processing and differential expression analysis

The raw sequencing files in FASTQ format were first aligned to Mus musculus genome (Ensembl 75) using STAR (version 2.3.0e) (44), and quantified by htseq-count (version 0.9.1) (45). Only genes with Counts Per Million (CPM) >=1 in more than six samples were retained for subsequent analysis. Batch effects were removed from the data using EdgeR (46). To capture astrocyte-specific signals, gene markers in other cell types (including neurons, microglia, oligodendrocyte, endothelial and ependymal cell), or genes highly expressed in other cell type compared against astrocytes, were further removed from the regional astrocyte signatures (47–50). Ribosomal genes curated in Gene Ontology were also removed to avoid ribosomal contamination.

#### Region-specific astrocyte transcriptional signatures

Regional astrocytes genes were determined as genes having significantly higher expression than other regions, calculated using EdgeR (46). As caudate nucleus and ventral striatum are similar in spatial location and gene expression profile (Figure 2A), gene signatures in these two regions were calculated without comparing with each other. P-values were subjected to Benjamini-Hochberg correction and region-specific astrocyte transcriptional signatures were defined as genes with FDR<=0.01 and mean CPM>=10 in that region.

#### Differential expression analysis in AD model

Differential gene expression analysis between AD and control mouse astrocytes were conducted using edgeR (46). Genes were ranked by their differential expression P-value.

### Construction of astrocyte-specific functional network

#### Gold-standards construction

To capture astrocyte-specific signal, we defined astrocyte-specific genes as the overlap of top quantile astrocyte-expressed genes in our TRAP dataset and highly expressed genes in astrocytes compared to other cell types (50). Known astrocyte markers identified from Human Protein Reference Database (HPRD) and other studies were also included (49). To ensure astrocyte specificity, we further removed gene markers of the other brain cell types, including neuron, microglia, oligodendrocyte, ependymal and endothelial cells (2,47,48). Mouse genes were mapped to human through ortholog mapping and functional knowledge transferring between species (51).

To capture functional relationships of each pair of genes in astrocytes, we first defined positive gold-standards as pairs of genes co-expressed and functionally related in astrocytes. Negative gold-standards include three types of relationships between pairs of genes: gene pairs not co-expressed in astrocytes, not functionally related, or neither co-expressed nor functionally related. The three types of negative gold-standards were randomly sampled to balanced ratio. Specifically, a pair of genes are defined as co-expressed in astrocytes when at least one gene belong to the astrocyte-specific genes in human, and the other gene is either an astrocyte-specific gene or a tissue-ubiquitous gene that is expressed in astrocytes (CPM>=10). A pair of genes are functionally related if they are co-annotated to the one of the expert-curated Gene Ontology biological processes (2).

#### Human brain functional genomics data compendia

Human brain data compendia were collected from 268 brain-associated gene expression datasets generated from 6,907 expression profiles, over 31,000 human protein-protein interactions, transcription factor binding profiles, chemical and genetic perturbation data as well as microRNA target profiles (23,2). For each dataset, we calculated gene-gene relationship scores based on the specific type of functional genomics measurements. Specifically, we downloaded brain-specific gene expression datasets from NCBI’s Gene Expression Omnibus (GEO) (52). Within each dataset, we calculated gene-gene coexpression as Pearson correlations between each pair of genes, applied Fisher transformation to the correlation coefficients and discretized the values into eight bins. Protein-protein interaction data were downloaded from BioGRID (53), IntAct (54), MINT (55) and MIPS (56).

Gene-gene relationships were calculated as binarized interaction scores between each pair of corresponding genes, except for BioGRID where interactions were discretized into five bins, ranging from no interaction to highest evidence of interaction. Based on motifs binding patterns in JASPAR database (57), gene-gene relationships were inferred from shared transcription factor regulation. For each gene, we scanned the 1kb sequence upstream for transcription factors binding (P<0.001) (58,59), and calculated scores between each pair of genes as Pearson correlation between the motif binding profiles. The correlation coefficients were further discretized into eight bins after Fisher transformation. We also downloaded chemical and genetic perturbation (c2:CGP) and microRNA target (c3:MIR) profiles from the Molecular Signatures Database (MSigDB) (60), and calculated relationships of gene pairs as the sum of shared perturbation profiles weighted by the specificity of each profile. The scores were further Fisher-transformed and discretized into eight bins.

#### Data integration

Using the gold-standards and pairwise gene relationships from the human data compendia, we trained a regularized Bayesian classifier to predict the functional relationship for each pair of genes in astrocytes (23). The Bayesian model that underlies our integration includes a class node indicating the presence or absence of a functional relationship between a pair of genes that is conditioned on hundreds of other nodes, one for each data set. The contribution of each data set is estimated in the model based on how relevant and accurate it is in reflecting how cellular pathways function in astrocytes. Since the assumption of conditional independence required for the naive Bayes classifier is violated for the large-scale genomics data sets, we incorporated regularization by calculating the mutual information between data sets and down-weighting similar data sets accordingly. The resulting astrocyte-specific network is a fully connected graph with 24,900 nodes as genes, and edge weights representing the likelihood of a gene pair participating in the same biological processes. The predicted posterior probabilities (i.e. edge weights) were scaled based on the assumption that the prior probability of a functional relationship is 0.01. Sleipnir library was used for the data integration (61).

### Network-guided regional astrocyte functional characterization

#### Region-specific astrocyte functional signatures and functions

For each astrocyte region, we used region-specific astrocyte transcriptional signature genes as queries in the astrocyte network to pull out genes specifically connected to them. To correct for the difference in network degree, for each gene in the genome, its connectivity to regional astrocyte queries was calculated as the fraction of its mean edge connectivity to the query genes compared to its mean edge connectivity to all genes in the network. More specifically, we calculate gene *i*’s connectivity to query set *Q* as 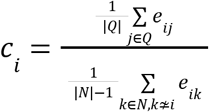, where *N* represents the collection of all nodes in the network, and *Q* represents the collection of region-specific astrocyte transcriptional signature genes, *e*_*ij*_ represents the network edge weight between gene *i* and *j*.

Region-specific astrocyte functions were then calculated by rank-based enrichment analysis on the ranked list of regional astrocyte functional neighbors with KEGG pathways, and pathways with FDR≤0.05 were retained.

### Network-guided reprioritization of disease differential expression analysis

#### Curation of astrocytic genes involved in Alzheimer’s pathogenesis

To compile a list of all astrocytic genes previously described to be associated with AD pathogenesis, an investigator completely independent from the analysis searched pubmed for English-language research articles with the keywords astrocytes and Alzheimer’s between 1975 and July 1^st^ 2018. Two lists were compiled: 1) an “expression” list: a list of genes described to be up-or down-regulated in AD brains, in the brains of AD mouse models, or in astrocyte cultures treated with Aβ, 2) a “functional” list: a list of genes functionally involved in AD. These genes were described to be, within astrocytes, functionally associated with AD pathogenesis, with amyloid load, with inflammation, with neuronal damage, with cognitive symptoms. If genes appeared in the expression and in the functional lists, they were automatically considered “functional” genes. When a publication described the involvement of a gene family in the astrocytic component of AD, but no experimental evidence was given to distinguish between family members, we did not include the gene family in our list.

#### NetDIFF algorithm to predict AD genes in astrocytes

We first defined positive gold-standards as significantly differentially expressed genes (P-value≤0.05, or top 1000 genes if there were less than 1000 significant genes meeting the stringent FDR cutoff), and negative gold-standards include genes with P-value>0.1 that are not within positive gold-standards.To ensure robustness, we subsampled one third of positives and negatives 50 times, with the probability of subsampling proportional to the differential expression P-value (positive samples are subsampled with probability of P-value^-0.2^, and negative samples are subsampled with differential expression P-value as probability). This enabled most significant genes being more likely to be chosen as positive and genes with minimal differential expression signal more likely to be selected as negative. A support vector machine classifier was then trained on each set of gold-standards to separate the positive and negative labels, using network connectivity in the astrocyte network as features. Finally, we aggregated 50 runs of predictions scores and generated an overall gene ranking on their likelihood to be perturbed in AD. To control for potential bias learned from the network structures, we randomized the differential expression P-values across genes 50 times, and performed support vector machine classifiers on the permuted gold-standards to generate a baseline prediction (background). Precision-recall curve was calculated on the predicted ranks in regard to the expert-curated gold-standards. Fold over background was then calculated by dividing the area under precision-recall curve (AUPRC) of NetDIFF prediction with the background baseline.

### Network-guided shortest path analysis to pinpoint mediator genes

#### Identification of the region-specific disease mediators (NetPATH)

To trace down the molecular path of how region-specific astrocytes may contribute to AD pathology, we leveraged the astrocyte network to identify genes connecting region-specific astrocyte transriptional signatures (regional signatures) to astrocyte-expressed AD risk genes (AD signatures). AD signaturess were defined as AD genes curated in OMIM and HGMD database with mean expression CPM≥10 in our TRAP dataset. To ensure computational efficiency and focus on strong functional connections, astrocyte network was filtered to retain edges with around top 1% of edge weights. Then, for each pair of regional signature and AD signature, we traced down the shortest path connecting in between them, with path length calculated as the sum of the reciprocal of each network edge weight along the path. Intuitively, for each gene in the astrocyte network, we calculated its betweenness centrality (BC) score as the fraction of shortest paths connecting the region-specific astrocyte transcriptional signature genes to AD genes through that gene. Thus, genes with high BC scores are likely molecular mediators between regional astrocytes and AD.

To ensure the genes identified are specifically mediating the connection between AD and regional astrocytes, we further performed permutations to rule out the scenario of identifying central genes in the network that have strong connections (i.e. on short paths) to most genes. To do that, we first computed each gene’s BC score connecting disease signatures with random genesets of the same size as regional signatures, and calculated a permutation-based P-value for each gene as the fraction of times that gene’s BC score with random disease geneset was equal to or greater than the gene’s BC score connecting the actual AD signatures (n = 100,000). Reversily, another P-value was calculated by replacing AD signatures with randomly generated genesets of same size as AD signatures 100,000 times. The two P-values were combined through meta-analysis and then subjected to

Benjamini-Hochberg correction. Genes were deemed important if they were on more than three shortest paths connecting regional signatures to AD signatures, and if the connection is weaker when using random regional and disease gene backgrounds (FDR≤0.1). To further ensure region-specificity, we performed this analysis on every region, and removed genes important between unrelated regions and AD from the list of region-specific disease mediators.

